# Movements for Grooming in Rhesus Macaques Co-opted from Feeding Movements

**DOI:** 10.64898/2026.10.26.761929

**Authors:** Ian Q Whishaw, Kristofers Buks, Angelina Ruiz, Vincent Savolainen, Amanda D. Melin

## Abstract

Catarrhine primates exhibit a wide range of skilled hand movements, including those used for foraging and grooming, and their evolution attracts ongoing attention. Competing hypotheses are that skilled hand movements evolved for feeding, or that the hand movements of various behaviors evolved independently. The present study addressed these questions by asking whether the movements of grooming of rhesus macaques (*Macaca mulatta*) are co-opted from feeding movements. We video-recorded free-ranging rhesus macaques in the Cayo Santiago colony of the Caribbean Primate Research Center (CPRC), University of Puerto Rico. Allogrooming (grooming a conspecific), self-grooming, and rhythmic scratching with a hand or foot were compared with a common foraging behavior, geophagy, mining and eating soil. Allogrooming, self-grooming, and foraging occurred in visually mediated bouts that featured numerous comb and brush hand movements directed toward searching for a target in the fur or soil. Once located, the target was grasped between the thumb and index finger in a pincer grip to be placed in the mouth - or less often bitted or licked from the fur. Hand movements were generally directed toward the actor but were also influenced by object-surface features, including fur direction during grooming and the rocks and debris covering soil targets during geophagy. The grooming movements of comb, brush, and grasp were recognizably those used during foraging. Rhythmic scratching with a hand or foot was not visually guided but did employ a comb movement of the digits of the hand or foot. Thus, with the exception of the rhythmicity of scratching, the hand movements of grooming are fundamentally foraging movements. It is proposed that grooming movements are coopted from oromanual feeding movements in macaques, a relationship that may be widespread in other species.

## Introduction

Skilled hand-shaping movements are used by vertebrate species for many behaviors, including locomotion, climbing, clinging, foraging and feeding, infant care, sexual behavior, aggression, play, social bonding, and body-surface care. In phylogenetic summaries of hand use, it has been proposed that the hand movements that evolved for feeding are also those used for many nonfeeding behaviors (Iwaniuk & Whishaw, 2000; Sustaita et al., 2013). Although this hypothesis has not been tested by comparing the hand movements used for different behaviors within a species, the detailed classifications of hand movements now available make such an investigation feasible. The hand skills used in eating have been classified into three groups: reach, grasp, and withdraw movements (Chehade & Gharbawie, 2023; Grant & Conway, 2019; Jeannerod, 1981; Jeannerod et al., 1995, 1998; Karl & Whishaw, 2013; Knecht, 2000; Marzke, 1971; Nashner et al., 1985; Sartori et al., 2015; Whishaw et al., 2019, 2025). Reach movements are hand movements made in relation to the extrinsic (spatial) properties of food items. They include power grips, in which objects are clamped by the fingers against the palm, open-hand shapes used to search for targets, and pointing movements. Grasp movements are made in relation to the intrinsic (size and shape) properties of food items. They include grips that secure a target between the distal thumb pad and one or more distal finger pads. Grips involving the thumb and index finger are termed pincer grips, whereas those involving the thumb and additional fingers are termed precision grips (Napier, 1956, 1962). Withdraw movements bring an item to the mouth, position it within the mouth, and hold it for successive placement into the mouth (An et al., 2026; Karl & Whishaw, 2019; Whishaw et al., 2025). The classification of hand use into these component acts is supported by evidence suggesting that each movement has its own evolutionary history. For example, animals such as rodents that do not ordinarily reach for food with a hand nevertheless use their hands to manipulate food after taking it into the mouth (An et al., 2026; Barrett et al., 2022; Whishaw et al., 2019). Strepsirrhine primates use visually guided power grips to pick up food but do not use precision grips and do not visually inspect food during withdraw-to-mouth movements, as do platyrrhine and catarrhine primates (Hirsch et al., 2022; Peckre et al., 2019, 2023; Whishaw et al., 2024). In addition, the relative use of grasp movements varies among platyrrhine primates. Capuchin monkeys use precision-like grasping strategies during feeding, whereas many other platyrrhine species rely on power grips (Christel & Fragaszy, 2000; Costello & Fragaszy, 1988; Pouydebat et al., 2008; Truppa et al., 2019; Whishaw et al., 2026). Accordingly, this research raises the question of the extent to which the diversity of hand skills displayed during feeding among different primate suborders and species is reflected in their other hand-use behaviors.

Given the proposed centrality of foraging movements to other forms of hand use, the present study examined the relationship between the hand movements used for eating and those used for grooming in rhesus monkeys. Grooming is a prominent and socially important behavior in rhesus monkeys and occupies a considerable proportion of their daily activity (Maestripieri and Hoffman, 2012; Spruijt et al., 1992). These monkeys use their hands for body-surface care, including parasite removal, cleaning, thermoregulation, stress reduction, and social bonding, behaviors that collectively comprise grooming (Dunbar, 1996; Sachs, 1988; Spruijt et al., 1992). Grooming includes allogrooming, in which animals groom conspecifics; various forms of self-grooming; and rhythmic scratching with the forelimbs and hind limbs. Despite numerous studies of the hygienic and social functions of grooming, descriptions of the movements themselves, with the exception of scratching, remain limited (Dunbar, 1996; Mooring, 2024; Sparks, 2017; Spruijt et al., 1992; Yerkes, 1933). Nevertheless, there are suggestions that grooming movements may have their own evolutionary history and, in turn, may even have contributed to the evolution of the hand skills used for feeding and foraging (Bishop, 1962; Dunkel, 2019; Mai et al., 2026; Spruijt et al., 1992; Goosen, 1981). Physiological and ethological studies of grooming in many vertebrate species further suggest that grooming movements consist of reflexes or stereotyped action patterns mediated by brainstem and spinal circuits that are distinct from feeding (Andronikashvili et al., 2024; Berridge et al., 1987; Berridge & Whishaw, 1992; Kalueff et al., 2016; Sherrington, 1906; Yang et al., 2021). For example, grooming using the forelimbs in rodents is organized and consist of stereotyped action sequences that can even be expressed in disorders affecting self-care (Berridge et al., 1987, 2021; Kalueff et al., 2016).

Because grooming incorporates both visually guided and stereotyped movements, it provides an excellent test of a key prediction of co-option, that feeding movements subserve grooming functions. Co-option proposes that movements evolved for one function can be subsequently used for different functions (Ding, 2025; McLennan, 2008). Allogrooming, self-grooming, and scratching were compared with a common foraging behaviour – geophagy (soil mining and eating) – in a free-ranging population of rhesus macaquesmacaques (*Macaca mulatta*) at the Cayo Santiago colony of the Caribbean Primate Research Center (CPRC), Puerto Rico. Geophagy was selected as the reference foraging behavior because this extractive behaviour is more representative of wild foraging activities than consumption of monkey chow, involves the search for edible items, and because the same monkey population that was videorecorded during grooming also engages in geophagy (Mahaney et al.,1995). The first aim of the study was to compare the reach, grasp, and withdraw movements used during grooming and eating and to assess their similarities. The second aim was to develop a classification of reach, grasp, and withdraw movements that could be applied in comparative studies of feeding and grooming in other primate species. To accomplish these aims, we conducted frame-by-frame analyses of video-recorded behavior, examined the syntactic organization of hand movements, and assessed the extent to which the movements were visually guided.

## Methods

### Study population

The results were obtained from video recordings 30 (14 female) adult rhesus macaques. The Cayo Santiago population of rhesus macaques (*Macaca mulatta*) was established in the late 1930s and started with 409 founders imported from India (Kessler et al, 2016). Since then, the macaques have flourished, with the population now approaching 2000 individuals. The population is managed by the Caribbean Primate Research Center (CPRC) at the University of Puerto Rico. The CPRC maintains a detailed census of the population, recording the date of birth of each monkey, and provisions them daily with monkey chow and water. Aside from tetanus inoculation administered when animals are approximately 1 year old, no medical interventions are administered to the individuals. The animals at Cayo Santiago are free to aggregate into social units and to supplement their diets with insects and leaves found on the island. They are exposed to natural weather conditions and interact in affiliative, agonistic, and reproductive behaviors with other members of the population. Each individual is recognized and tracked by tattoos, ear notches, and facial features. Interventions are minimal. The research was approved according to the Institutional Animal Care and Use Committee of the Caribbean Primate Research Center (CPRC) at the University of Puerto Rico, permit IACUC A500118.

### Video recording

Video recording at 30 frames per sec (fps) using a VIXIA HF G70 camcorder or using an iphone 11 provided ad libitum recordings of natural foraging and grooming behavior. The behavioral results were obtained from frame-by-frame analysis of QuickTime Player (v. 10.5), and frame counts were obtained from the frame count option of the playback.

### Behaviors studied

Figure 1 illustrates examples of the behaviors that were studied in which three types of grooming were compared with geophagy.

**Figure 1.**
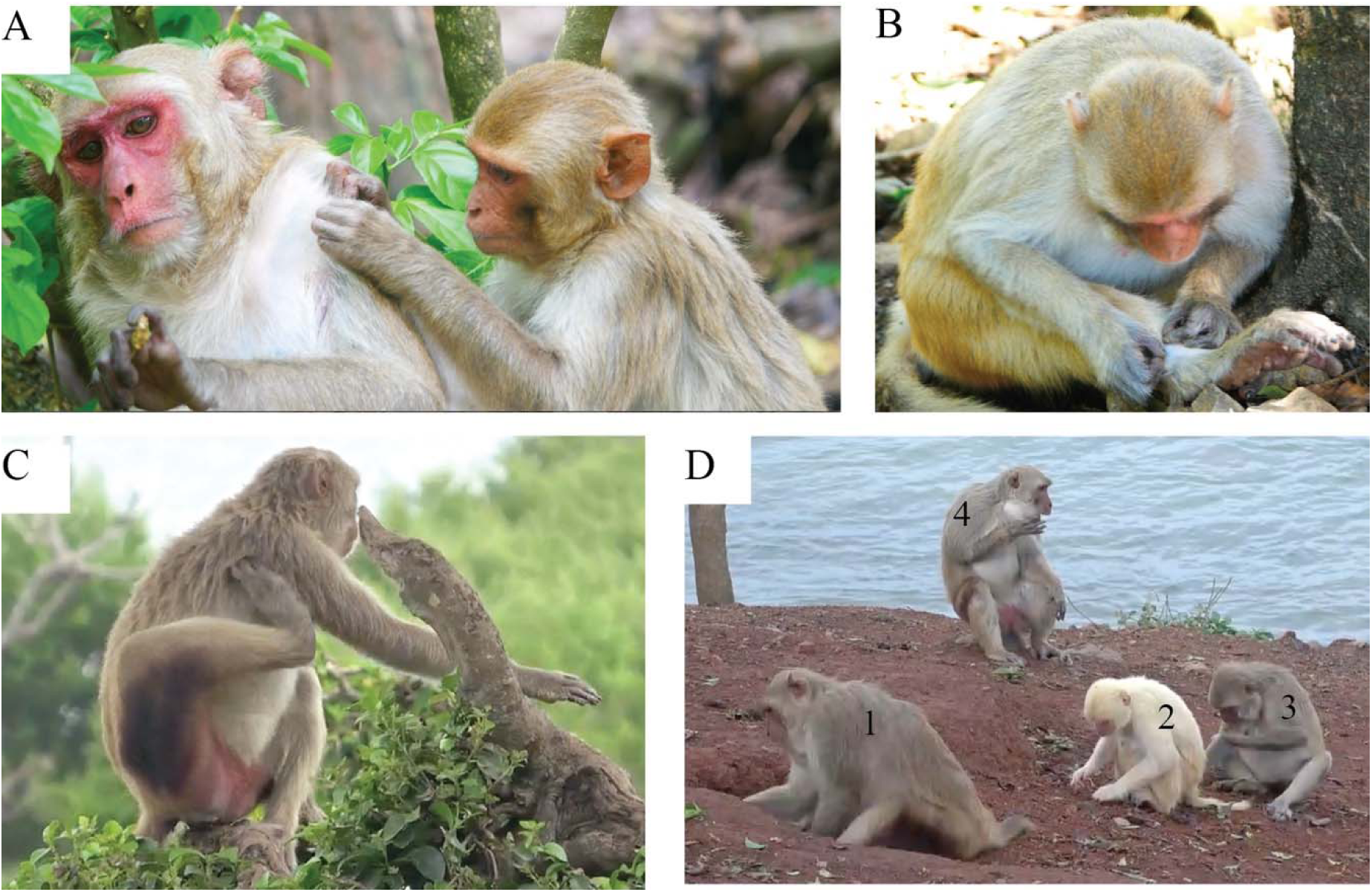
Hand use associate with four different behaviors in Rhesus macaques. Note: A. Allogrooming using both hands by an adult female rhesus monkey directed to the back of an adult male monkey. B. Self-groom by an adult female with both hands to examine its right lower leg. C. Scratching using the right hind foot directed to the upper arm. D. Geophagy by a group of rhesus monkey: (1) mining, (2) grasping soil with a left hand pincer grasp, (3) withdrawing soil to the mouth to eat using the left hand, (4) adjusting soil with the right hand in the cheek pouch.

A. *Allogrooming*. Allogrooming (n=10 monkeys, 5 female) movements were hand or mouth movements directed toward another monkey and included observations on adults grooming each other or infants.

B. *Self-grooming*. Self-grooming (n=10 monkeys, 4 female) movements were hand or mouth movements that a monkey direct toward its own body surface

C. *Geophagy*. Soil eating (n=10 monkeys, 5 female) movements included: (1) digging into the soil with the hands, (2) brushing rocks and debris from the ground surface, (3) grasping soil and placing it in the mouth, and (4) adjusting consumed soil in the cheek pouch.

D. *Rhythmical Scratching*. Rhythmical scratching movements were scratching movements of a hand or foot directed toward scratching a relatively focal part of an animal’s own body and these were obtained from most of the monkeys as they engaged in the other behaviors.

### Behavioral bouts

A behavioral bout was defined by the duration of gaze engagement for the behaviors of allogrooming, self-grooming and geophagy. Bout duration was obtained by converting frame counts to seconds. A bout onset was scored as beginning when a monkey turned its gaze fully toward the location of hand movements on a target. A bout offset was scored when a monkey oriented its head to look in another direction or brought its hand to its mouth, which usually involved a concomitant head raise often accompanied by a blink (Hirsch, 2022).

### Hand and mouth movements

Preliminary observations of the rhesus macaques engaged in allogrooming and self-grooming and geophagy showed that each bout consisted of a number of hand and mouth movements and these were described and counted for the present study.

1. *Hand movement*. Hand movements were defined as any discrete hand contact made by a monkey with a target and included movements of bringing a hand to the mouth. Counts of all hand movements were made for each bout.
2. *Hand movement direction*. The direction of a hand movement during contact with a target was designated using an Eshkol-Wachman numeric-derived system (Golani, 1994; Whishaw et al. 2024). In reference to the tip of the snout, “0” represents a movement of the hand directly away from the snout and 1-7 representing 45^0^ steps of clockwise rotation. For example, a hand movement that brought the hand to the mouth was scored as “4”, representing a movement directed toward the snout. The movements of the left and right hands hand were scored separately.
3. *Mouth grasp movements*. Each mouth movement that consisted of advancing the mouth to a target on the fur was counted. If the tongue visibly licked associated with a mouth movement, the mouth movement was scored as a lick.
4. *Kinematic representation.* Kinematic measures of head and hand movements were made by digitizing body points using the program Tracker (v 6.4.3).

### Positional behavior

The posture adopted by and an animal when allogrooming, self-grooming, scratching, or soil-eating was recorded as sitting on the haunches or standing on all four limbs.

## Statistical Analyses

Gaze duration, hand grasping type, laterality are reported as total number of observations for allogrooming, self-grooming, geophagy or scratching. Conjunctions of behaviors were assessed using a Pearson product-moment correlation. Group comparisons (e.g., gaze duration, hand use, behavior, etc.) were made using a generalized linear model repeated measure analysis of variance using SPSS (29.0.1.1) or t-tests that designated probability and power. Follow up tests were Newman-Keuls or Bonferroni tests. Group numeric comparisons were made with a chi-square (χ^2^) test and variance was assessed with Levene’s test. The temporal relationship between movements were described kinematically. Statistical values of p<0.5 were considered significant (Cote et al., 2021).

## Results

### Behavioral Bouts are Associated with Visual Gaze

Figure 2 illustrates kinematic representations of gaze and hand movement associated with a representative allogrooming bout by a monkey that is grooming a conspecific (Figure 2A). The kinematic representation of the mouth (tip of the upper lip) and the left and right hands (the knuckle of the second digit) illustrate movement of the groomer during one bout (Figure 2B). The bout begins with the face orienting to the target and ends with the face disengaging the target. Eight grooming movements are made by the left (the hand is open and brushes the fur of the conspecific) and right hand (the hand is semi-closed and combs the fur of the conspecific) during the bout. The arrows on the recipient monkey in Figure 2A indicate the location and direction of the hand movements made by the groomer, including the hand to mouth movement.

**Figure 2.**
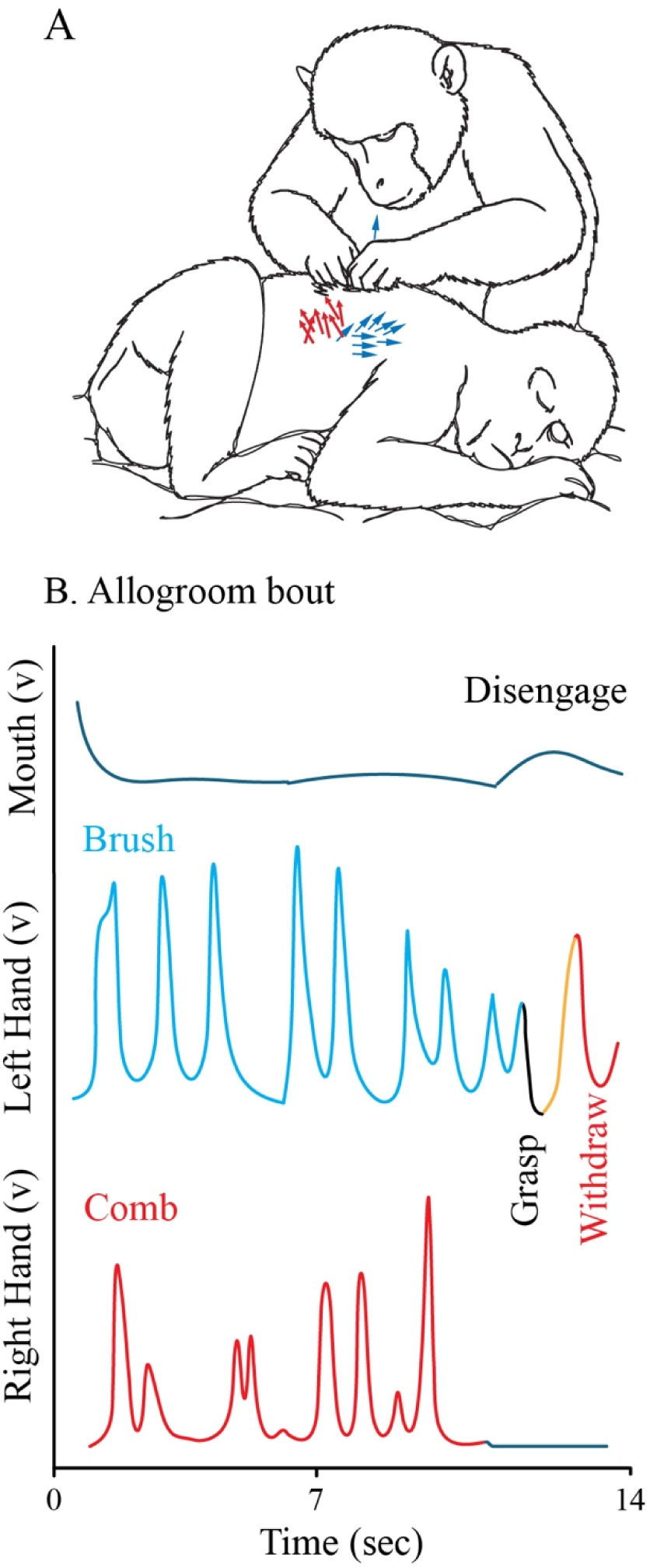
A kinematic representation shows the movement of the head and right and left hands during an allogroom bout. Note: A. Direction of hand movements made by the left and right hand of the groomer. B. The groom bout lasts 14 seconds starting with a head movement to direct gaze to the grooming target and ending with a gaze disengage as the head raise to take a grasped target into the mouth from the left hand. The left hand makes 8 brush-movements across the fur followed by a grasp and withdraw-to-mouth of an item found in the fur. The right hand makes 8 comb-movements to part the hair.

### Bout duration and hand movement number related

Figure 3 illustrates (A) allogrooming, (B) self-grooming and (C) geophagy. For all three categories, no monkey was observed to make hand movements contacting a target in the absence of directed visual gaze toward the target. Sometimes a hand movement directed to a target began just before gaze was engaged and sometimes a hand movement finished an act just after gaze was disengaged, and these movements were also counted as being associated with a gaze event.

**Figure 3.**
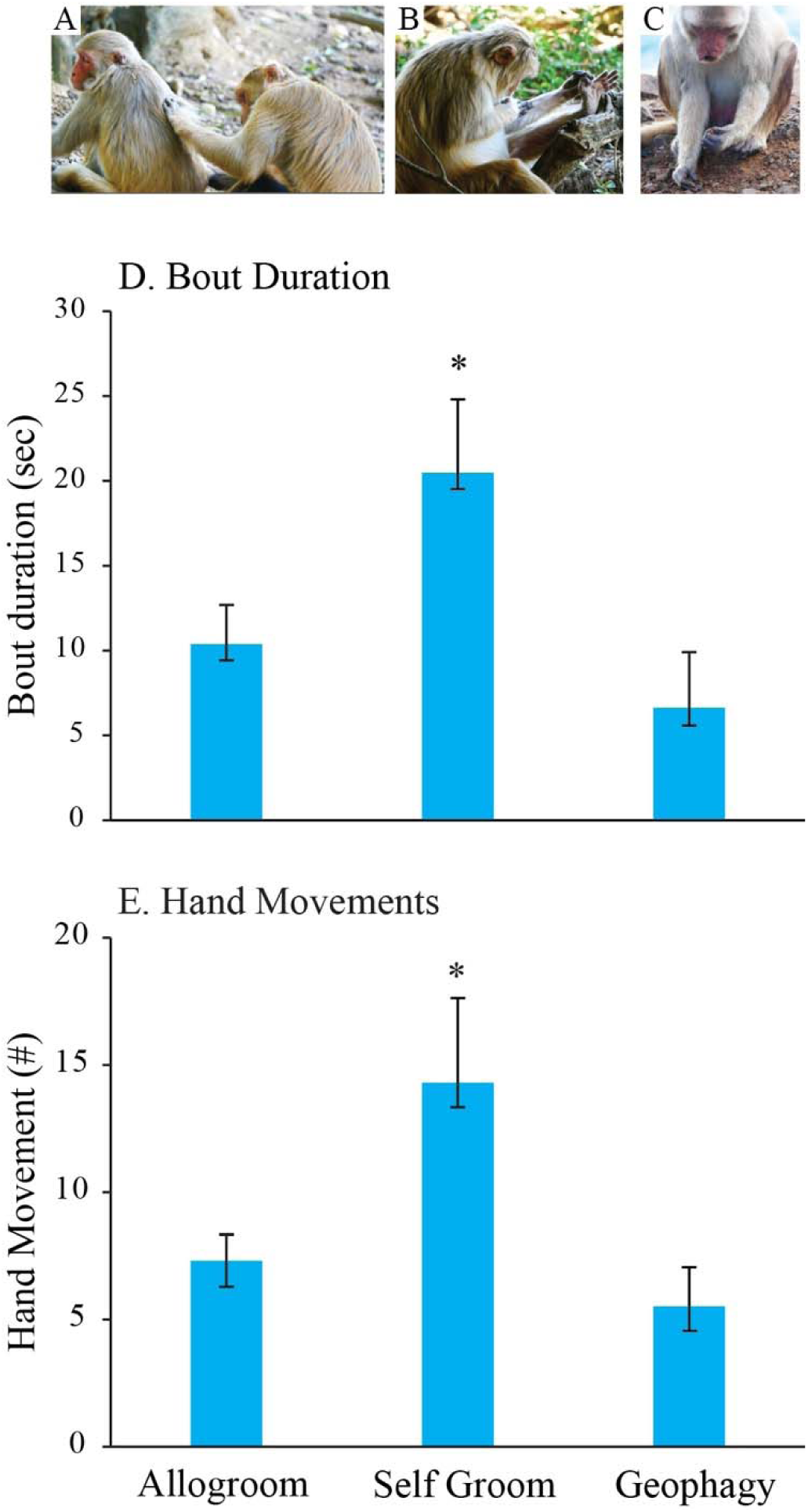
Bout duration associate with allogroom, self-groom and geophagy movements. A. Allogroom, B. Self-groom, C. Geophagy. D. Mean±se of bout duration associated with allogroom, self-groom and geophagy behavior. Self-groom durations are significantly longer (* p<.05) than allogroom and geophagy durations which do not differ.

Figure 3D gives the mean±se of duration of behavioral bouts for two types of grooming and for geophagy. Figure 3E gives the mean±se of hand movements associated with a bout. In all, 364 bouts (88 allogrooming, 98 self-grooming and 178 geophagy) were measured. We found a significant effect of Duration, (one-way ANOVA, F(2,29) = 5.83, p=.008), and Hand Movement Number (F(2,29) = 4.50, p =.021). Follow-up Newman-Keuls tests showed that for allogrooming vs geophagy, differences in duration (p =.380) and number (p =.790) were not different. Duration was longer for self-grooming vs allogrooming (p =.003) and self-grooming vs dirt-eating (p =.031). Number of hand movements was larger for self-grooming vs allogrooming (p =.009) and self-grooming vs soil-eating (p =.013). Similar long durations of self-grooming is reported by (Boccia, 1983).

Figure 4 summarizes durations and hand movement number as log/log scattergrams and correlations for all bout durations for all monkeys. Correlations were: (A) allogrooming (r =.62, p <.001) (B) self-grooming (r =.81, p =.001) and (C) geophagy (r =.59, p <.001). Although significant, the correlations are not high because there was scatter between gaze duration and the number of hand movements. For example, an animal may make a single grasp and withdraw to the mouth with a gaze duration lasting less than a second or make a single hand movement across the fur or the dirt and then stare without further hand movements for many seconds.

**Figure 4.**
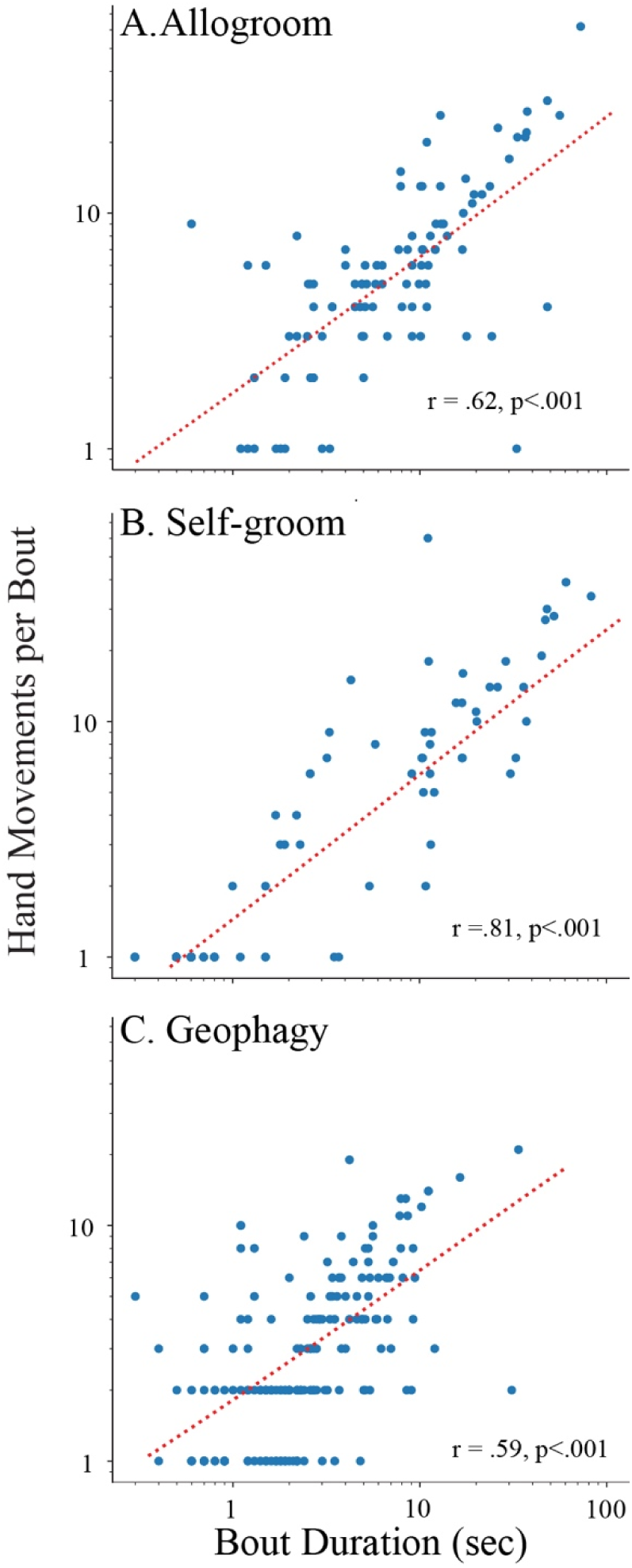
Log-log plots of the hand movement number as a function of bout duration for allogroom, self-groom, and geophagy. Note. Significient correlations featured in each scatter plot indicate that the execution of hand movements during each behavior accounts for most of the duration of the bouts associated with each behavior.

### Hand movements similar in grooming and dirt-eating

Figure 5 illustrates three types of hand movements that occurred during allogrooming, self-grooming and soil eating.

**Figure 5.**
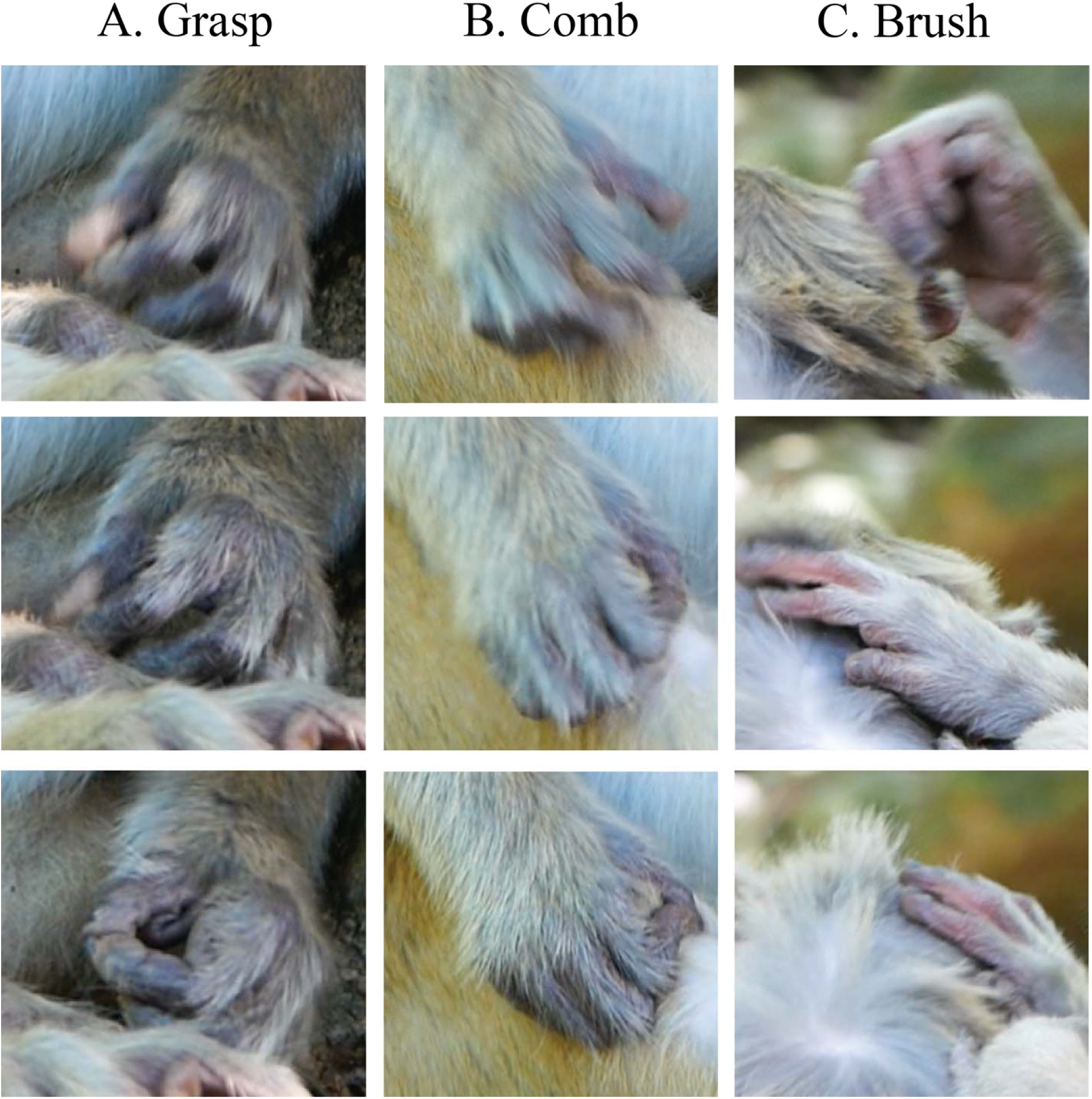
Hand movements associated with allogooming. Note. A. Grasp. The hand is pre-shaped in anticipation of the grasp (top), makes the grasp (middle) and grasps the target with a pincer grasp between the pads of the thumb and first finger. B. Comb. The hand is pre-shaped in anticipation of the grasp (top), the fingers are flexed to initiate the comb (middle), and the finger tips comb the fur as the hand is pulled toward the body (bottom). C. The hand is preshaped (top), opened as the hand contacts the fur (middle) and the hand brushes the fur with the palmar surface of the fingers and palm (bottom).

### Grasp

A pincer grasp movement is a hand movement that grasped a target held between the thenar pad and the pad of the index finger oriented in opposition. Figure 5A shows three phases of the grasp, in which the digits are flexed as they preshape for the grasp (top), closed to make the grasp (middle) and then oriented the hand by wrist rotation (bottom) for visual examination of the target or to bring the target to the mouth. Every grasp observed for allogrooming, self-grooming and soil-eating was a pincer grasp. A partial exception was one monkey engaged in allogrooming who had a seeming injury to the index finger and so made a pincer grasp using the thumb opposed to the middle finger. Grasp movement could be followed by a monkey holding the presumptive grasped target up for visual inspection to bring it, with a withdraw of the hand, to the mouth. Grasp movement could occur singly or repeatedly as a monkey might pick and pull at a target.

### Comb

Comb movement were hand movement in which the fingers were flexed as they were dragged across the target. Figure 5B shows three phases of a comb movement. The hand was preshaped with the fingers flexed before contact (top), contacted the fur with the fingers flexed (middle) and dragged across the fur while further closing (bottom). Comb movement could involve all of the fingertips, a variable number of fingertips or one fingertip in contact with the target. Comb movements resembled power or whole hand grasping movements in that all the fingers but not the thumb were used.

### Brush

Brush movements were made with the palmar surface of an open hand. Figure 5C shows three phases of a brush movement. The fingers lightly flexed in anticipation of contact (top), opened to make palmar contact (middle) and moved across the surface of the fur (bottom). In a brushing movement the entire palmar surface, the radial or the ulnar side of the palm or only the surface of a single digit might be in contact with the target.

### Incidence of grasp, comb and brush movements

Figure 6 illustrates as percent the numbers of grasp, comb and brush movements associated with allogrooming, self-grooming and geophagy (actual numbers of movements are also given). When allogrooming, brush movements were most frequent, when self-grooming grasp movement were most frequent and during geophagy, the numbers of each type of movement were similar, yielding a significant Chi-Square (4) =176.4, p<.001. An ANOVA for repeated measures gave no differences in the incidence of left vs right hand use for the different behaviors, F(1,27) = 1.28, p =.266.

**Figure 6.**
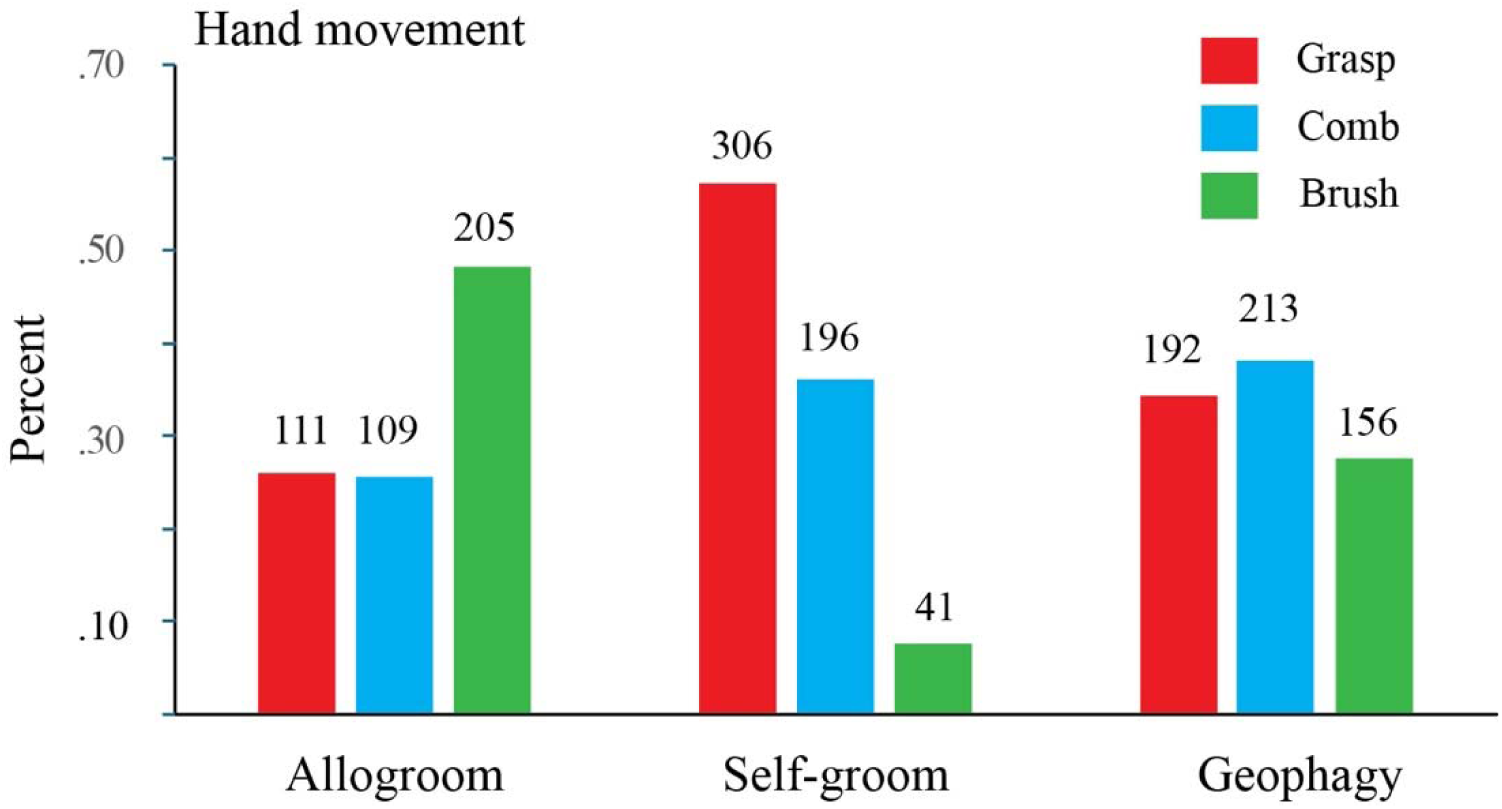
Distribution of hand movement types for allogroom, self-groom and geophagy as a percent. Note. Allogroom is associated with more brushes and self-groom more grasps, whereas the distribution of hand movements in soil-eat is similar.

**Figure 7.**
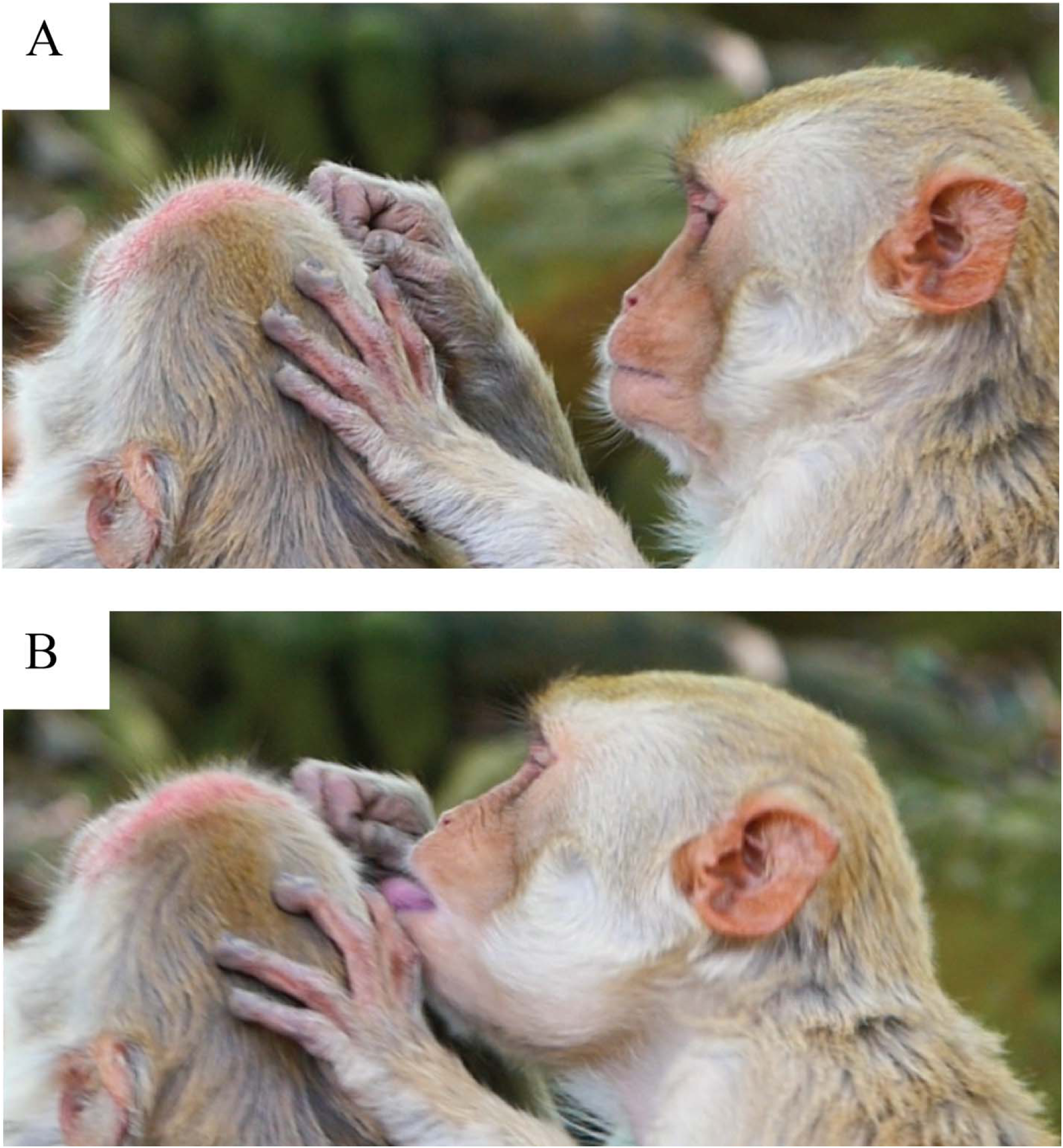
Tongue use in allogrooming Note: The left hand as brushed the fur to the left and the right hand has combed the fur to the right and then the rhesus monkey advances its mouth and protrudes its tongue to lick the fur.

### Mouth grasp movements also used for allogrooming

Mouth grasp movements were also used to take objects from the fur when allogrooming, but mouth grasping was not observed during self-grooming or soil-eating. Figure 6 illustrates a mouth grasp in which the tongue is used. In Figure 6A (Video 1) the monkey makes made a brush movement with the left hand and a comb movement with the right hand and has apparently visually identified a target on the fur. In Figure 6B, the monkey advances its head and protrudes its tongue to take the target from the fur with its tongue. In all, 61 mouth grasps were observed by the allogrooming monkeys, with all from 3 monkeys that had 61, 8 and 1 mouth grasp. Of these, 15 mouth grasps were observed to involve tongue grasping (11, 3, 1). In should be noted, however, that once the mouth contacted the fur, it was difficult to determine how the mouth was used for grasping, and so it can only be concluded that the monkeys sometimes did use their tongue to lick and at other times used their mouth to grasp. The monkey that made the most mouth grasps was only observed to make only mouth grasps, whereas the other monkeys more frequently made hand grasps.

### Hand movement made with lower arm flexor movements

Figure 8 gives polar plots of the direction of hand movements in allogrooming, self-grooming and soil-eating. A within-subjects ANOVA gave a significant effect of hand Direction, F(1,26) = 41.09, p<.001, with most hand movements made inward toward the monkey or laterally to its sides (e.g., pulling movements), with very few movements directed away (e.g., pushing movements). The comparison of hand direction when allogrooming, self-grooming and soil-eat, were not significantly different, F(2,26) =.519. p =.601. Because lateral hand movements frequently made laterally (direction 2 &3 for the right hand and direction 5 & 6 for the left hand), left hand directions were flipped in order to make a composite hand use comparison, which gave no significant difference in how the hands were use, Hand F(1,26) = 2.04, p =.164.

**Figure 8.**
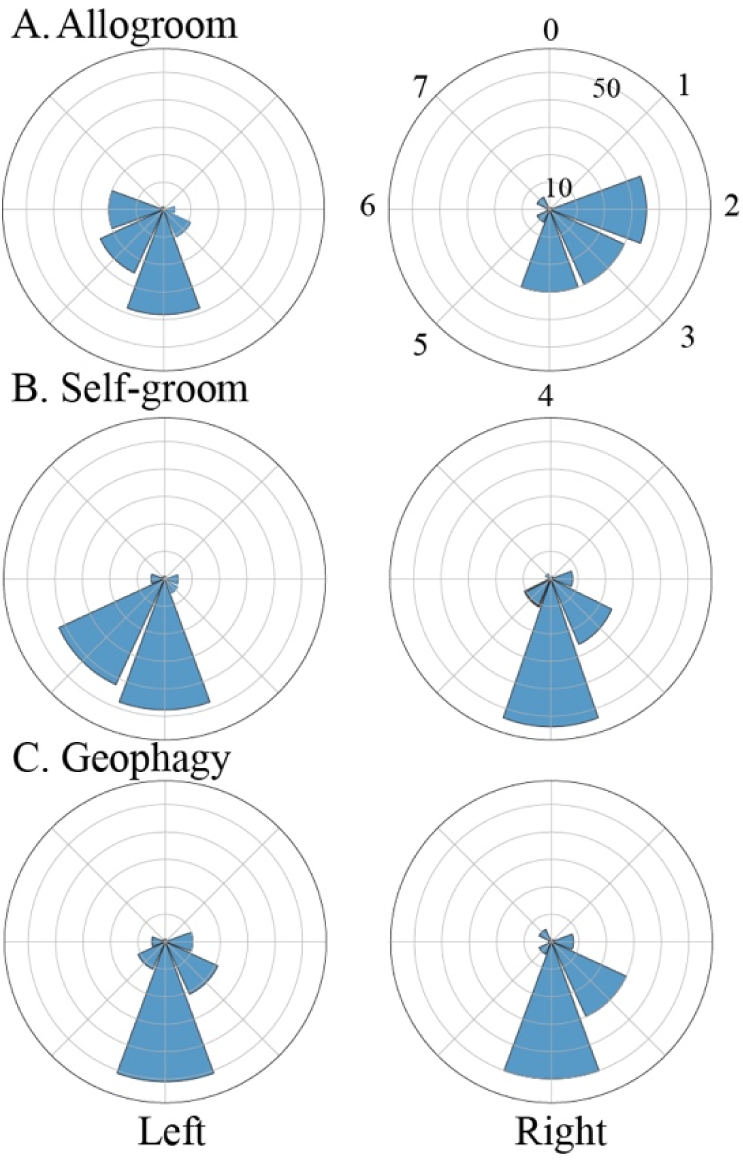
Polar plots of the direction of hand movements for allogroom, self-groom and geophagy Note. Each polar plot represents the horizontal orientation of the body with the face pointing toward zero. The plots show that most hand movements are directed toward the body, i.e., a result of a flexion movement of the arm. Grasp hand movements that resulted in the hand bringing an object to the mouth are made in the 4-direction.

### Scratching made with the forelimbs or hindlimbs

Figure 9 summarizes the analysis of scratching using the left or right forelimb or hind limb, in which scratches were variously directed to the back, head, arms and legs. Figure 9A illustrates a forelimb scratch and Figure 9B (Video 3) illustrates a hind limb scratch. In all 17 forelimb scratch events featuring a total of 247 scratches (average = 14.5 scratches per event) and 8 hind limb scratching events totalling 136 scratches (17.5 scratches per event) were observed. For none of the scratching events was gaze directed toward the target that was scratched. The hand movements for scratching were very similar to comb movements, in which the digits were advanced and opened toward the target of the scratch, closed (with more closure seen for the forelimb scratches) as contact and the scratch occurred. Figure 9D shows scatter plots and correlations for scratch duration and the number of scratches per event, which gave a significant correlation for the forelimb scratches (r =.75, p<.001) and the hindlimb scratches (r =.96, <.001). The mean±se frequency for forelimb (.294±.086) and hindlimb (.341±.094) were not significantly different, t (23) = 1.07, p =.296. A Levene’s test on group variance also produced a nonsignificant result for forelimb scratching and hindlimb scratching, F=.163, p =.214.

**Figure 9.**
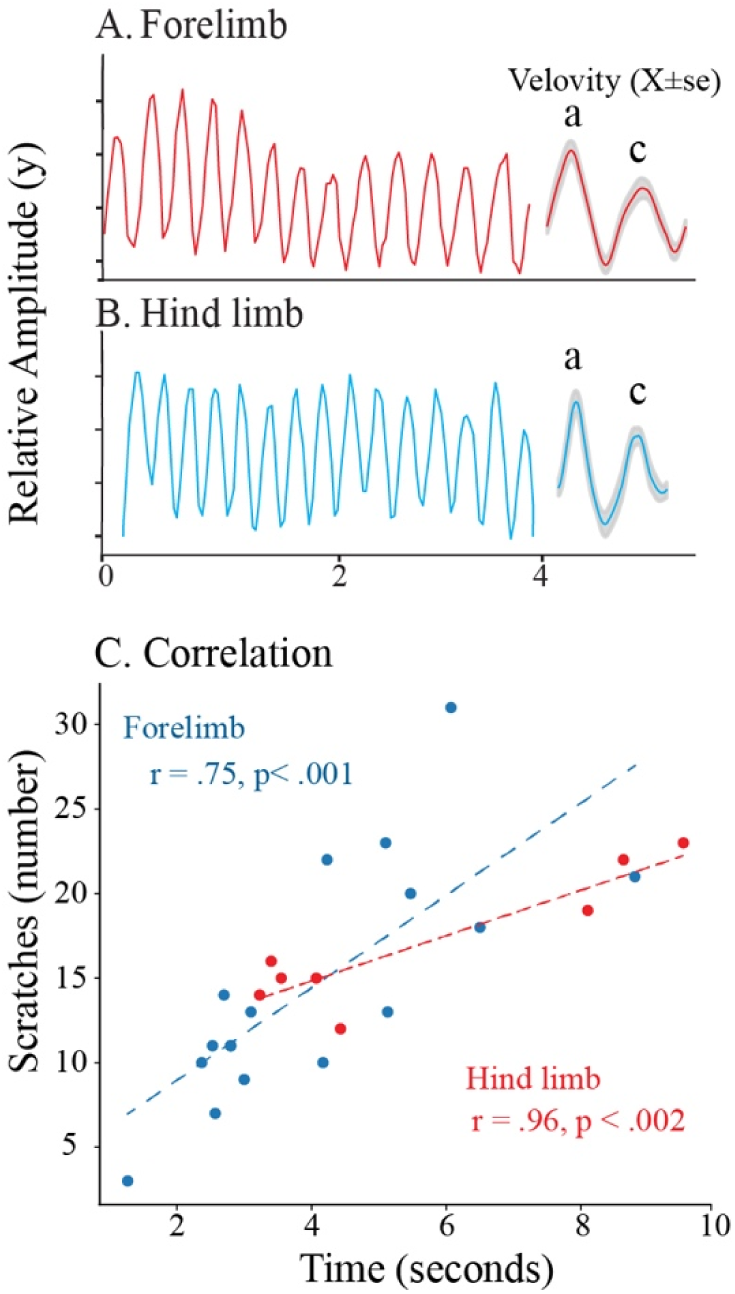
Forelimb and hind limb rhythmical scratching. Note: A. Kinematic examples of forelimb and hind limb rhythmical scratching (y-axis). The insert shows average velocity in which the first peak represents the limb being lifted to scratch and the second peak representing the downward movement of the scratch. B. Significant correlations of the number of scratches in a scratch bout as a function of time for both forelimb and hind limb. a, advance, c, comb.

## Discussion

The aim of this study was to investigate the hypothesis that skilled hand movements associated with foraging and grooming share an evolutionary origin. We find that the hand movements used by rhesus macaques for grooming are highly similar to the hand movements they use for eating, providing support for the hypothesis that they are co-opted from feeding. During all occurrences of allogrooming, self-grooming, and geophagy, hand movements occurred in visually guided bouts. The hand movements used during grooming consisted of two types of reach movements (brush the fur with an open hand and comb through it with flexed fingers) and one grasp movement (a precision grip) and a withdraw movement to bring items to the mouth. Precision grips picked items from the fur, held them up for inspection and placed them in the mouth for eating. The monkeys used the same brush and comb movements to remove debris covering the soil, after which soil particles were also taken with a pincer grip for geophagy. Unlike grooming, scratching with either the forelimb or hind limb was rhythmic and not visually guided, but the scratch itself was produced by digit flexion similar to that of a comb movement. Thus, with the exception of the rhythmic component of scratching, the results show that grooming movements closely resemble feeding movements. These findings are consistent with a conclusion that feeding movements are co-opted for self-grooming and allogrooming movements.

The co-option of feeding movements for grooming movements may be a general rule. Figure 10 presents a model that outlines the idea as well as the idea that the evolution of skill oromanual behavior will provide the skills for other behavior. Available evidence suggests that co-option of feeding movements of other behavior is widespread (Held, 2017; Hodgson et al., 2024). Grooming in mice is characterized by highly stereotyped action patterns, nevertheless examination of the forelimb movements of their grooming reveal that they resemble those used during eating (Naghizadeh et al., 2020). Although there has been no detailed analysis of grooming in strepsirrhine primates, several observations indicate that although they may use their hands to part the fur during allogrooming, they do not display a hand withdraw but use the mouth to remove objects from the fur (Dunkel, 2019; Buettner-Janusch & Andrew, 1962; Spruijt et al., 1992). This parallels their feeding, in which they hold food in a hand but reach with the mouth to retrieve it (Peckre et al., 2023). For platyrrhine monkeys, howler monkeys do not use the hand to grasp and withdraw food to the mouth (Whishaw et al., 2026) and anecdotal reports suggest that they display relatively little hand allogrooming (Chiarello, 1995). For platyrrhine monkeys, direct corticospinal projections are absent in squirrel monkeys but are present in their close relatives, capuchin monkeys (Bortoff & Strick, 1993; Strick et al, 221). Correspondingly, squirrel monkeys do not use precision grips, whereas capuchin monkeys do, suggesting that their grooming should be correspondingly different (Christel & Fragaszy, 2000; Spinozzi et al., 2004). Although not examined here, it is possible that the motivation and reward associated with grooming are similarly derived from foraging, as there is evidence for overlap in their brain networks (Berntson1973; Micco 1974; Spruijt et al, 1992). We cannot rule out that evolutionary selection for skilled grooming could further refine skilled foraging, and this idea could be subject to investigation (Dunkel, 2019).

**Figure 10.**
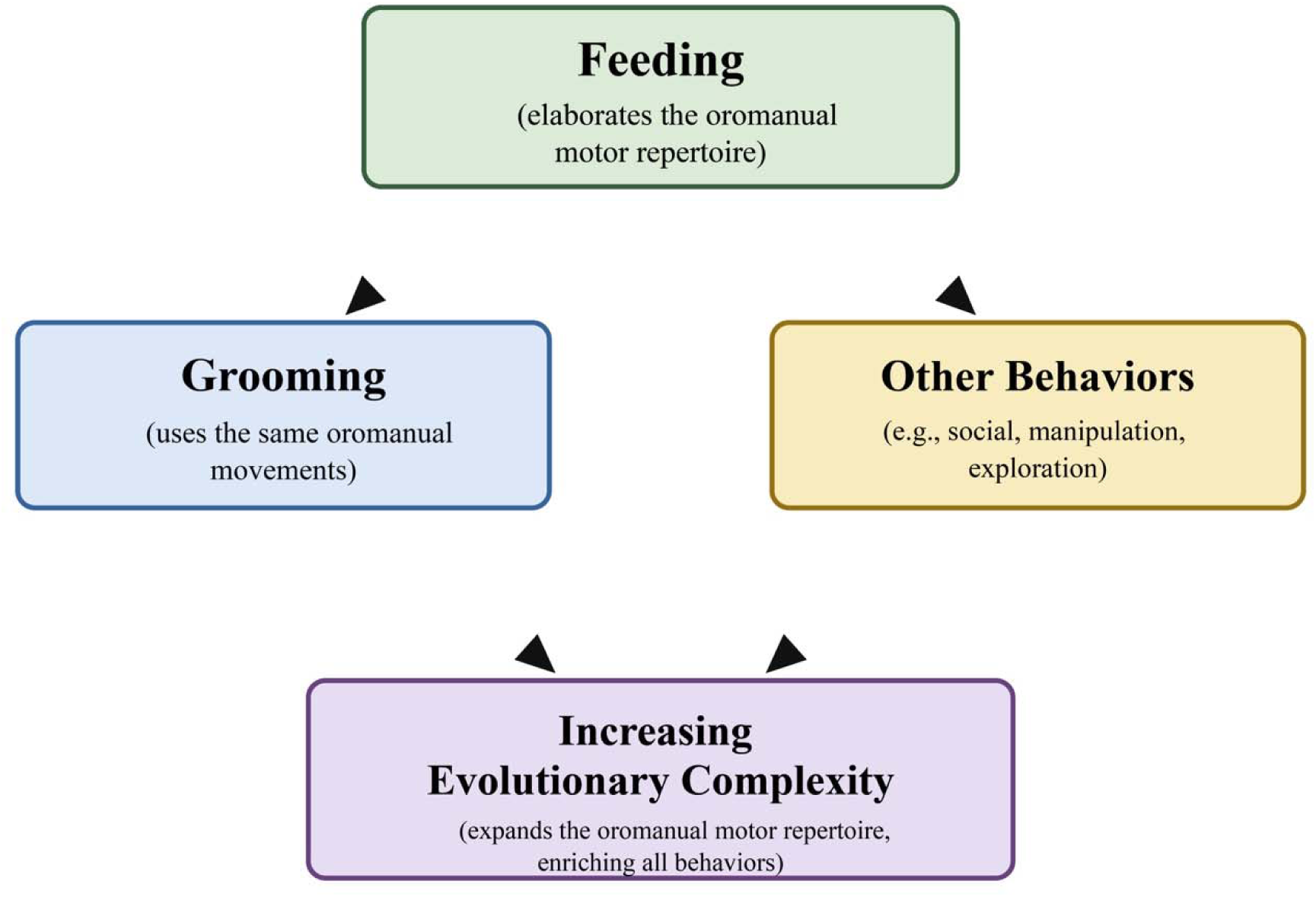
Theoretical model of the relationship between feeding and grooming. Note: The model illustrates the oromanual repertoire of skilled movements is co-opted for grooming and likely for other behaviors and proposes that evolutionary acquired feeding skills will similarly be reflected in grooming and other behavior.

The hand shapes identified during allogrooming, self-grooming, and geophagy are consistent with the proposal that catarrhine primates employ three broad categories of hand movement: reach, grasp and withdraw (Grant & Conway, 2019; Jeannerod, 1981; Jeannerod et al., 1995, 1998; Karl & Whishaw, 2013; Knecht, 2000; Marzke, 1971; Napier, 1956, 1962; Nashner et al., 1985; Sartori et al., 2015; Whishaw et al., 2019, 2025). The reach movements of brush and comb manipulated the fur during grooming and removed debris during the soil mining components of geophagy. They served a search function, consistent with the proposal that reach movements primarily subserve spatial function. The open-hand search movement resembles that used by blindfolded humans searching for a target (Karl et al., 2012) and the open-hand posture when reaching into far peripheral vision, where the precise location of an object is uncertain (Hall et al., 2014). It is unlikely that the reach movements of brush and comb are restricted to geophagy behavior described here. In our video archive, we have observed rhesus macaques foraging in trees using similar brush and comb movements to part leaves while searching for edible items. These activities have not been recorded frequently enough to permit formal analyses. The reach hand movements displayed subtle variations when used during grooming and soil eating, and these differences appeared to reflect the physical properties of the substrate being manipulated. For example, brush movements for geophagy appeared to be faster than those used during grooming, and comb movements were used for digging. Despite these differences, the appearance and direction of the movements toward the body were remarkably similar during both grooming and feeding. A pincer grip used to withdraw objects to the mouth was distinctive and was the only grasp movement used to remove debris from the fur during grooming or to grasp soil particles for inspection and eating. This grip is clearly suited to the small size of the objects manipulated during both behaviors. Considerable evidence suggests that the pincer grip is supported by direct corticospinal projections onto spinal motor neurons in catarrhine primates (Lemon, 2004, 2019), and the present results extends the conclusion that it mediates precision grasping for grooming in addition to feeding.

Not only were the hand movements used during feeding similar to those used during grooming, but the overall organization of the two behaviors was remarkably similar. Grooming occurred in visually guided bouts that began with gaze directed toward a grooming location and ended when a target was brought to the mouth - or by interruption by another monkey. A gaze bout could last for only about a second, during which a monkey made a single reach to grasp an item from the fur and bring it to the mouth, or could continue for more than a minute, accompanied by repeated hand movements. Thus, it was evident that both grooming and feeding involved sustained visual attention. Eye tracking methods would be useful for measuring gaze duration, eye blinks (Hirsch et al., 2022) and attentional saccades (Kobayashi & Hashiya, 2011), because the monkeys’ heads were usually directed downward their eyes were not sufficiently visible to make these distinctions. For humans, interblink intervals during tasks requiring sustained visual attention, such as reading (Bentivoglio et al., 1997), may approach 10 sec, approximately the average duration of the gaze bouts observed during grooming and geophagy in the present study.

The stimuli that elicit rhythmic scratching and the neural circuits that support it have been extensively investigated in a number of species, including cats, dogs, and turtles (Murphey et al., 2025; Sherrington, 1906; Stein, 2018). In contrast, scratching in primates has often been regarded either as an abnormal displacement behavior or simply as another component of self-grooming and has received relatively little attention (Diezinger & Anderson, 1986; Mootnick et al., 2012). Nevertheless, Mootnick and colleagues present phylogenetic evidence suggesting that rhythmic forelimb scratching appears early in primate evolution, being present in strepsirrhines and platyrrhines, and becomes the exclusive form of scratching in apes. It is doubtful that rhythmic, whole-arm scratching occurs in humans, in whom nonvisual brush and comb movements of the fingers are more typically described as responses to itching (Ayres, 1964; MacKay-Lyons, 2002). The present study suggests that rhythmic scratching with the forelimbs and hind limbs is similar in rhesus macaques with respect to scratching frequency and movement pattern. Furthermore, the digit shaping of both the hands and feet is a reach movement, as the fingers or toes flex but do not grasp. Thus, although scratching shares digit configurations with the combing movements used during grooming and feeding, its rhythmic organization distinguishes it from visually guided grooming behavior.

There has been considerable research on the social functions of grooming in rhesus macaques, including investigations of sex differences, social relationships, and hand preferences (Beisner et al., 2026; Boccia et al., 1982; Dunbar, 1996; Fu et al., 2023; Sachs, 1988; Spruijt et al., 1992). Although grooming by both sexes and by monkeys of different ages was observed in the present study, the experimental design was not intended to examine these aspects of grooming behavior. A more comprehensive identification of individual animals and substantially larger sample sizes would be required to address these questions. Instead, the present study focused on the types of hand shaping used during grooming and feeding. We found no evidence that age, sex, or individual behavioral differences influenced the patterns of hand use that we observed. Nevertheless, future studies examining the development of hand shaping and the effects of aging on grooming and feeding movements in rhesus macaques would be valuable. We also observed one monkey featuring an injured index finger used a precision grip that was formed between the thumb and the distal pad of the middle finger and we also observed that one monkey only used mouth grasping for targets on the fur, suggesting that rhesus macaques display considerable flexibility in the organization of skilled hand movements.

In conclusion, rhesus macaques use a common repertoire of structured reach, grasp and withdraw hand-shaping movements during grooming that resemble the hand movements used during feeding. These findings support the hypothesis that species-typical feeding movements have been co-opted for other hand-related behaviors. This conclusion further suggests that the motivational and reward systems underlying grooming may recruit neural mechanisms that originally evolved to support feeding behavior. For both grooming and feeding, an important component of motivation may be the search for a target suitable for ingestion, with reward ultimately achieved by placing that target into the mouth. We propose that the behavioral organization of grooming and feeding described here provides a comparative framework for examining the evolution of hand use in other primate suborders and species.

## Acknowledgements

We thank former and current staff at the Caribbean Primate Research Centre for support, in particular Francelia Marte, Giselle Caraballo, Bianca Giura, Josue Negron, Daniel Phillips, Nahiri Rivera, Carmen Ramos, and Imperial student Clarence Toh for help with data collection.

## Author Contributions

IQW: Conceptualization, Methodology, Formal analysis, Investigation,Resources, Data curation, Writing-Original Draft, Writing-Review & Editing, Visualization,Supervision, Project administration;

KB: Videorecording, Writing-Review& Editing

AR: Investigation, Writing-Review& Editing;

VS: Investigation, Writing-Review& Editing;

ADM: Methodology, Data curation,Writing-Review & Editing, Supervision, Project administration, Funding acquisition.

## Videos

**Video 1**.

Allogroom by a female rhesus monkey of a male rhesus monkey in which the left hand makes a brush movement, the right hand makes a comb movement followed by a grasp, and the female monkey then uses her tongue to remove a target on the fur.

**Video 2**.

Geophagy by a female rhesus monkey in which the left hand makes two brush movements followed by a series of comb movements followed by a pincer grasp of some soil using the right hand.

**Video 3**.

A series of 12 scratches to the upper chest with the right hind foot. On the initiation of each scratch the toes are extended and then, during the movement of the limb, flex to make the scratch.

